# A point cloud segmentation framework for image-based spatial transcriptomics

**DOI:** 10.1101/2023.12.01.569528

**Authors:** Thomas Defard, Hugo Laporte, Mallick Ayan, Soulier Juliette, Sandra Curras-Alonso, Christian Weber, Florian Massip, José-Arturo Londoño-Vallejo, Charles Fouillade, Florian Mueller, Thomas Walter

## Abstract

Recent progress in image-based spatial RNA profiling enables to spatially resolve tens to hundreds of distinct RNA species with high spatial resolution. It hence presents new avenues for comprehending tissue organization. In this context, the ability to assign detected RNA transcripts to individual cells is crucial for downstream analyses, such as in-situ cell type calling. Yet, accurate cell segmentation can be challenging in tissue data, in particular in the absence of a high-quality membrane marker. To address this issue, we introduce ComSeg, a segmentation algorithm that operates directly on single RNA positions and that does not come with implicit or explicit priors on cell shape. ComSeg is thus applicable in complex tissues with arbitrary cell shapes. Through comprehensive evaluations on simulated datasets, we show that ComSeg outperforms existing state-of-the-art methods for in-situ single-cell RNA profiling and cell type calling. On experimental data, our method also demonstrates proficiency in estimating RNA profiles that align with established scRNA-seq datasets. Importantly, ComSeg exhibits a particular efficiency in handling complex tissue, positioning it as a valuable tool for the community.

## Introduction

Understanding the spatial organization of tissues at the single-cell level is crucial to study disease and development (1–3). Molecular profiling of single cells in their spatial context allows to infer cell states and cell types, cell-cell interactions and cell-fate decision-making (4), as well as the study of the overall tissue architecture, under normal and diseased conditions, leading to the definition of spatial domains and disease signatures (5).

Spatial transcriptomics (ST) denotes a large panel of technologies that allow the spatially resolved measurement of gene expression. These methods can largely be divided into two groups: sequencing-based (6, 7) and imaging-based (8–10). The former measure the entire transcriptome, but has a lower spatial resolution, while the latter relies on measuring a subset of genes. Such a subset consists of pre-defined marker genes that are specific for the cell states or types of interest. Since not the entire transcriptome is imaged, we will refer to these approaches as RNA profiling. These imaging-based approaches are variants of single-molecule fluorescence in situ hybridization (smFISH) methodologies and can detect RNAs with single-molecule sensitivity with high spatial resolution, several orders below the scale of a single cell. A typical dataset consists of 2D or 3D point clouds of the imaged RNA species. One challenge in the analysis of these data is the correct assignment of these RNAs to their cells of origin. Indeed, in contrast to single-cell RNA sequencing (scRNA-seq), the information on which RNA molecules belong to the same cell has to be inferred from the image. This assignment is crucial, as it impacts cell type identification and thus the major aspect of the analysis we would like to perform.

One frequently used approach to perform this assignment is to segment the cells from additional channels employing markers for cell and nucleus segmentation. RNAs are then assigned to the cells based on their spatial positions with respect to this segmentation. Such stainings typically encompass labeling of the nucleus with DAPI, cellular staining with one or more cell membrane dyes or labeling all RNAs as a proxy for the cytoplasm (10–12). However, cell membrane staining is often not an option (13, 14). Besides, staining can be technically complex (15), and may not work equally well for the entire tissue, thus leading to inhomogeneous cell segmentation and cell type calling accuracy. Further, the boundaries of individual cells can be challenging to segment, especially for tissue with complex 3D cellular morphologies.

More recently, several computational approaches have been presented to detect individual cells in the images and establish their RNA profile without relying on a dedicated cell marker. These methods rely only on the RNA positions and in some cases a DAPI stain, to regroup RNAs according to local transcription profiles. Such RNA point clouds can then act as an approximation for cell shape, by establishing a hull that encompasses all RNAs that were deemed to belong together (15–17). While these approaches hold great promise for the analysis of spatial RNA profiling data, their use is limited by implicit priors on cell shape (15, leading to underperformance in complex tissues or requirement of additional scRNA-seq data (16, 17) (seeTable 1).

**Table 1:** Characteristics of existing methods for spatial RNA profiling.

|  | Watershed | pciSeq<br>(16) | JSTA<br>(17) | SSAM<br>(18) | Baysor<br>(15) | ComSeg |
| --- | --- | --- | --- | --- | --- | --- |
| Based on convex shape prior | Yes | Yes | Yes | No | Yes | No |
| Require external dataset | No | Yes | Yes | No | No | No |
| Single-cell profiling | Yes | Yes | Yes | No | Yes | Yes |

While there exist already methods for spatial RNA profiling, these methods usually come with requirements that are not always met in practice. For instance, some methods require a high RNA density which often implies a large panel of marker genes. However, while current commercial spatial RNA profiling approaches provide hundreds of marker genes, they remain extremely costly, and for many questions, fewer marker genes will be sufficient, which can be probed with simpler custom-built solutions (19–21). Furthermore, most methods implicitly assume convex or even round cell shapes. In contrast, the cell shapes in some tissues can be complex and often deviate from such simple shapes. Approaches relying on strong assumptions on cell shape are therefore suboptimal for many of these tissue types. Lastly, some methods require parallel scRNA-seq data, which in practice is not always readily available, so ideally the use of such information should be only optional.

To alleviate these issues, we propose our new method ComSeg. ComSeg uses as input only the coordinates of the RNA molecules and optional spatial landmarks, such as nuclei. ComSeg then groups RNAs with similar expression profiles, optionally aided by these landmarks. It does not necessitate scRNA-seq data or cytoplasmic markers, nor does it make implicit use of any prior assumptions regarding cell morphology. Instead, the method describes RNA point clouds as graphs weighted by gene co-expression and relies on graph community detection. Our method is easy to apply by design as it does not require complex machine learning model training. Furthermore, we provide the tool as an open-source Python package (https://github.com/tdefa/ComSeg) with extensive documentation: https://comseg.readthedocs.io, compatible with the scverse environment (22).

Development of such analysis approaches requires annotated ground truth to assess their performance. Experimental ground truth can be obtained in some cases by employing membrane markers, from which the cytoplasmic membrane might be segmented with deep neural networks or manual annotation (23). However, the ground truth quality is affected by the heterogeneity of the staining quality which is low on complex and dense tissue (15). Furthermore, experimental data does not permit a more systematic exploration of parameters such as the expression level or cell morphology. Hence, similarly (16, 17), we address the lack of high-quality ground truth by generating simulated data allowing us to control the complexity of the input data. We develop *SimTissue (https://github.com/tdefa/SimTissue)*, an open-source Python simulation package to reproduce in-silico fluorescent-based spatial transcriptomic experiments.

We used the *SimTissue* framework to validate ComSeg on simulated data with increasing complexity in terms of abundance of RNA, number of marker genes, and tissue morphology. ComSeg outperforms other methods in terms of Jaccard index for RNA-cell association in most of the tested scenarios. We also validate ComSeg on an in-house experimental dataset of mouse lung tissue imaged with smFISH and human embryonic lung tissue (24) imaged with HyBISS (25). On these experimental data, ComSeg estimates accurate RNA profiles that match established scRNA-seq datasets. Overall, the shape agnostic approach of ComSeg demonstrates notable efficacy for complex tissue composed of cells with non-convex shapes.

## Material and Methods

### Comseg Overview

ComSeg associates the detected RNA molecules with their corresponding nucleus, cell landmarks or cell centroid. It leverages a k-nearest neighbor (KNN) graph, where the nodes are the detected RNAs. The method can be decomposed into 5 steps:

1. Computation of a proximity weighted expression matrix
2. Construction of KNN graph weighted by co-expression
3. Graph community detection optionally using landmarks, such as nuclei or cells
4. Cell segmentation free in situ clustering of communities
5. Final RNA assignment

The only hyper-parameters exposed to the user are the mean cell diameter *D* and the maximum cell radius *R*_*max*_. The numerical values of these hyper-parameters are detailed in the supplementary methods (section 1).

#### 1) Proximity weighted expression matrix

ComSeg leverages gene co-expression information. In principle, co-expression information can originate from parallel scRNAseq data or from other published resources. However, such external data sets are not always available for the biological system that is studied. Hence, we estimate co-expression using only the spatial arrangements of detected RNA molecules in the image. For this, we leverage the spatial correlation between the different RNA species molecules as a proxy for gene co-expression.

##### Spatial correlation computation

For each RNA molecule *m*_*x*_ at *x*, we compute a proximity weighted expression vector (PE) *V(x)* summarizing the local RNA profile of the molecule *m*_*x*_ by considering its *K*= 40 closest neighbor molecules *y* ∈ *KNN(x)* in a radius *R*_*pe*_=D/ 2.

*V(x)* is defined as the sum of neighbors *y*, weighted by a score, that linearly decreases between the position of *x* and the circle around *x* with radius *R*_*pe*_. To compute *V(x)* we first compute *V*_*g*_ *(x)* for each marker gene *g:*

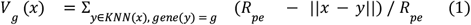

By calculating *V*_g_ *(x)* for all marker genes, we get the PE vector *V(x)* defined as:

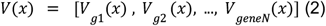

By stacking the *V(x)* for each molecule *m*_*x*_, we get a proximity weighted expression matrix 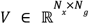 where *N*_*x*_ is the number of detected RNA and *N*_*g*_ the number of marker genes.

From this, we can finally compute the co-expression matrix 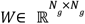, where each element *w*_*i,j*_ is the Pearson correlation of the columns of *V* corresponding to the expression of genes *i* and *j*:

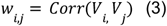

The computation of these co-expression values is implemented by the Python class *ComSegDataset* of our package. Of note, the co-expressions could alternatively be computed from external data, e.g. from single-cell RNA sequencing data.

#### 2) KNN graph weighted by co-expression

Our algorithm operates on a weighted KNN graph, where the RNA molecules (across all genes) are the nodes (*K* = 10 and the max edge distance *R*_*knn*_ between molecules is set to *D*/ 4) and the weights are the estimated co-expression values *w*_*i,j*_ defined in (3): edges between RNA molecules from strongly co-expressed genes obtain a large weight, while RNA molecules from genes that are not co-expressed are assigned a small weight. The rationale is that RNAs from co-expressed genes are likely to belong to the same cell, while RNA molecules from genes that are usually not expressed together are more likely to belong to different cells.

#### 3) Graph community detection algorithm

In the previous section, we generated a graph strongly connecting RNA nodes that are likely to belong to the same cell. Now, our objective is to partition this graph into sets of RNAs belonging to the same cells. To achieve this, we developed a modified version of the Louvain method (26) for community detection so it can accommodate prior knowledge given by nuclei segmentation or other landmarks.

The original Louvain algorithm is a widely used method to partition a graph into sets of strongly connected nodes. These strongly connected sets of nodes are called communities. The algorithm optimizes a metric of graph structure called the modularity, noted *Q*(27). *Q* is the sum of the differences between intra-community weights and their expected value in a randomly rewired graph. *Q* can be expressed as a sum over the edges (*u,v*) of the graph.

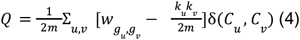

Where *g*_*u*_ and *g*_*v*_ are the gene index of the RNA nodes *u* and 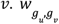 is the corresponding weight from the the co-expression matrix *W* .*k*_*u*_ is the degree of node *u* defined as 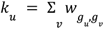 is the sum of the weights in the network 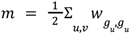 and the function δ is 1 if the nodes *u* and *v* belong to the same community *C* (i.e *C* _*u*_ = *C* _*v*_) and 0 otherwise.

This method iterates two elementary phases: in the first step, modularity is greedily *Q* optimized. For this, we start from an initialization where each node is assigned to its own community. Nodes are then moved to neighboring communities to greedily maximize modularity. This first step stops when no node move improves the modularity *Q*.

In the second step, a new aggregated network is built where communities found in the first step become nodes. Edge weights between those new nodes are calculated as the sum of the edge weights between the identified communities. We can then re-apply the first step on this aggregated network until there is no modularity gain.

When applying the Louvain method, we only consider positively weighted edges as negatively co-expressed genes are not likely to belong to the same cell. Besides, in order to introduce prior knowledge in the form of landmark segmentation if available, we modify the method as follows: before running the community detection method, RNA nodes inside the same segmented nucleus or chosen cell landmark are merged together into one node, which we refer to as “cell nodes”. When we apply the Louvain method, different cell nodes cannot be merged together. During the first step of local moving of nodes, nodes take the cell label of the community they are assigned to.

Of note, in most cases, nuclear staining is available, and this therefore represents the most frequent use case. However, the landmarks can also originate from other stainings (such as membrane staining) or the analysis of RNA density or any other method that would allow to identify a subset of RNAs that clearly belongs to the same cell.

The input graph strongly connects RNAs that are likely to belong to the same cell. Hence the resulting community partitions of RNAs are supposed to form sets of RNAs belonging to only one cell. In contrast, a cell might contain several communities.

The graph construction and partitioning are implemented in the Python class *ComSegGraph* of our Python package.

#### 4) Cell segmentation free in situ clustering

In the previous step, we split our graph into communities of RNAs that are supposed to belong to the same cell. Hence, while a cell can be composed of several RNA communities, we assume that each community does not extend beyond cytoplasmic boundaries. This mimics the mechanism of superpixel segmentation in computer vision (28).

To ease the identification of RNA communities that may belong to the same cell, we first identify communities that have similar RNA profiles.

We associate with each community *C* of RNA, an expression vector *V*_*C*_ .

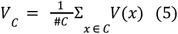

where *V* (*x*) are proximity weighted expression vectors (PE) computed as defined above. Summing the PE of each RNA node in the community helps to capture the local transcriptomic information missing in the global co-expression matrix. Hence each *W* community is associated with a community expression vector *V*_*C*_ composed of co-expressed genes at the global scale but also containing the local transcriptomic information of the cell it belongs to.

Then, similarly to what is commonly done in scRNA-seq analysis (29), we cluster our set of community expression vectors *V*_*C*_ using optionally PCA for dimensionality reduction (depending on the number of marker genes) and the modularity-based algorithm Leiden (30). It defines community clusters that exhibit similar expression profiles. Each community (and each member *m*_*x*_) thus receives a community profile label *L*_*i*_. However, we do not cluster community expression vectors of less than three RNAs as they might be too small to reliably capture their local transcriptomic neighborhood. We assign to these small communities the majority label of the nearest neighbors.

This step thus provides a map of RNAs labeled with their community profile label *L*_*i*_. We refer to this map of labeled RNAs as the transcriptomic domain map. The in-situ clustering step is implemented in the class *InSituClustering* of our Python package.

### Final RNA assignment

In the final step, the goal is to associate the RNAs with the cell they belong to. First, we add a centroid node, a node that does not correspond to an RNA molecule. If the landmarks contain nuclear staining, this centroid node is the centroid of the nucleus, all nuclear RNAs are merged into this node, and the centroid node gets the community profile *L*_*i*_ of the nuclear RNAs. In case there is no nuclear RNA, the community profile of the centroid is defined as the majority profile among the *K* nearest neighbors (*K* = 15, *R*= *D* / 2). In case there is no nuclear staining, the centroid node can be defined based on other landmarks (e.g. the maximum of the distance function of a cellular landmark) or the analysis of RNA density. While cell or nuclei landmarks are optional, these cell centroid nodes are required to estimate the single-cell RNA profiles.

Once every cell centroid is associated with a community profile *L*_*i*_, we can finally estimate the single-cell RNA profiles. We associate each cell centroid of label *L*_*i*_ with their nearest RNAs of the same label. We employ the geodesic distances between cell centroids and RNA nodes. The geodesic distance is the graph’s shortest path distance between the centroids and RNA nodes and is computed with the Dijkstra algorithm (31)

. We apply the Dijkstra algorithm with Euclidean distance weight on the edges. Besides, when an RNA node’s geodesic distance to its nearest centroid is superior to a chosen maximum cell radius *R*_*max*_, the RNA molecule is not associated with any cell. Using the geodesic distance helps to better estimate the cell size of non-convex cells when applying the cell radius *R*_*max*_ cut-off. Moreover, geodesic distance accommodates potential lacunar spaces within the tissue when calculating the centroid-RNA distance.

In summary, our method treats RNA positions as nodes in a graph with edges weighted by co-expression. This permits an accurate separation of cells with different expression profiles without the need for explicit cell segmentation. Our method can further use landmarks, such as nuclei to initiate the community detection in spatial RNA graphs. In the absence of clear differences in expression profiles, cells will then be separated based on RNA-centroid distance in the graph.

Our method does not use an explicit cell shape prior other than mean cell diameter and *D* maximal cell radius *R*_*max*_ as the goal is to make this method suitable for tissues harboring arbitrary cell shapes. Furthermore, ComSeg is not based on Machine Learning, and does therefore not need annotated data or time-consuming learning steps; which simplifies its application and interpretability.

## Simulation

### Simulation Python package

In order to validate our method and perform quantitative benchmarks against other approaches, we designed a simulation framework *SimTissue* capable of creating ground-truth data with tunable complexity for tissue morphology, cell type compositions, and marker-gene expression levels.

Our simulation framework can be divided into two steps:

1. Simulation of tissue morphology.
2. Simulation of RNA composition and spatial distribution.

*SimTissue* is implemented in Python and available at https://github.com/tdefa/SimTissue.

#### Simulation of tissue morphology

Our framework offers two possible simulation scenarios. In the first scenario, we consider regular geometric patterns. While these are not realistic scenarios, they are well suited to point to potential problems and limitations of the algorithms. They can thus be seen as a purely methodological test scenario. Examples include the checkerboard arrangement or the simulation of clamped L-shapes with random nuclei positions.

In the second scenario, we consider more realistic tissue simulations. For this, *SimTissue* takes as input segmented nuclei from experimental data. These segmentation masks can often be generated easily from DAPI or other nuclei stainings and are widely used for image-based spatial transcriptomic experiments (8, 13, 32). Individual cytoplasms are defined by growing cells from segmented nuclei. Each cell grows at a random speed to add irregularity to the cell size. Still, some organs, such as lung tissue, contain lacunar space without cells. Therefore, optional masks can be added to indicate these lacunar spaces where cells cannot grow into.

In experimental data, the tissue section is cut at an arbitrary location and some nuclei are removed from the rest of the cell. It is hence possible to simulate cells without nuclei to better mimic experimental fluorescent-based experiments.

#### Simulation of RNA composition and spatial distribution

Once the nuclei positions and cell shapes are simulated, we have to simulate the RNA composition and distribution within each cell. RNA expression levels can either be set as constant or be drawn from a provided experimentally measured distribution, e.g. from scRNA-seq data. When drawing a profile from scRNA-seq, we sample for each simulated cell an expression profile of a single cell from scRNA-seq, then multiply the number of RNAs by a factor. Here, we choose a factor of 3 as the fraction of mRNA transcripts captured per cell in scRNA-seq data can reach 30% (depending on reagent chemistry) (33) while the capture rate of smFISH experiment is close to 100% (34). Finally, RNA molecules are randomly positioned in the available space of the cell with a uniform spatial distribution.

In summary, *SimTissue* allows to simulate experiments of incremental complexity. The full control over the ground truth and the difficulty of the segmentation task aim to understand the limitations of the benchmarked methods.

### Simulations in this article

#### Regular pattern simulations

We simulated 2D square grids (15μm x 15μm) and nested L-shape patterns (four squares of 15μm x 15μm). The pixel size is 0.150μm and thus similar to the pixel size obtained with a 60x objective. Nuclei are spheres of 3.75μm rays. Nuclei are in the center of the cell for square cell shape. For L-shaped cell, the nucleus is randomly positioned in the center of one of the four squares composing the L-shaped cell.

#### Lung tissue simulations

We simulated 3D mouse lung tissue leveraging experimental FISH data from (32). The original data is composed of images of 112μm x 150μm in XY and 15μm in Z with a DAPI staining and a Cy3 fluorescent channel. The original pixel size is 0.103×0.103 μm and the Z spacing is 0.300μm. The positions of the nuclei in our simulation are the positions of the segmented nuclei in the original images. The nuclei were segmented with Cellpose (35) on DAPI staining.

Next, we identified the space occupied by cells by thresholding the Cy3 FISH signal (first quintile of the Cy3 distribution). Individual cytoplasms were defined by growing cells with random irregular speed from segmented nuclei inside the allowed space. Random growth speed allows to add irregularity in the cell size. Finally, we removed 20% of nuclei in our simulation to simulate nuclei missed during sample preparation, as explained above.

We used a list of 34 cell type marker genes with the objective to classify 19 different cell types present in mouse lung tissue. The list of marker genes was selected using the NS-forest algorithm (36) on our external scRNA-seq dataset of mouse lung tissue (32). Then, we associated each individual cytoplasm with an expression vector sampled from our scRNA-seq dataset so that all the RNA profiles in our simulation are taken from real experimental data. This dataset can be found at https://zenodo.org/records/10172316.

### Benchmarking of ComSeg against state-of-the-art methods

We benchmark ComSeg against pciSeq (16), Baysor (15) and Watershed method on both simulation and experimental data. These are among the frequently cited and most widely used methods today if membrane markers are absent. The hyper-parameter setting of the benchmarked methods can be found in the Supplementary Methods.

#### Benchmarking on simulations

To assess the RNA profiling quality we compute the mean Jaccard index as follows:

For each cell 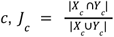 where *X*_*c*_ is the ground truth set of RNAs associated with the cell *c* (ground truth) and *Y*_*c*_ is the set of RNAs predicted as associated with the cell *c*. The final mean Jaccard index per cell is:

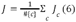

For each cell *c*, we compute the percentage of wrongly associated RNA *WA*_*c*_ as follows (False Discovery Rate):

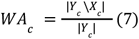

We compute the percentage of missing RNA *MS*_*c*_ (False Negative Rate) per cell as follows:

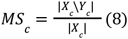

To perform cell type calling from the cell expression vector from Baysor, Watershed and Comseg, we first normalize the count matrix from both scRNA-seq and from RNA-cell association using the scTransform normalization (37). Then we compute the cosine distance between the cell expression vector and the cell type median centroid defined in the reference scRNA-seq data from (32). Cells are classified into their nearest cell type cluster in terms of cosine distance. For lung tissue simulation, we also employ this cell type calling method on the ground truth expression vector of each cell to assess the maximum accuracy achievable with the 34 selected markers.

Converly to previously cited methods, pciSeq performs cell type classification and RNA-nuclei association simultaneously. For this reason, the cell type classification described here was not applied to pciSeq.

### Experimental evaluation

We applied the benchmarked methods on two datasets of lung tissue. The first one exhibits 6 different marker genes in 3D mouse lung tissue acquired with a home-built sequential smFISH system. This mouse lung tissue was irradiated with 17 gy five months prior to mouse sacrifice as described in (32). The second dataset was acquired with HybISS (25), consisting of a human embryonic lung and 147 genes in 2D (24). In both cases, we perform a nucleus segmentation with Cellpose (35) and use this segmentation as initialization in our benchmark.

### Reference scRNA-seq cluster

As we do not have ground truth for experimental data, we leverage scRNA-seq to check the consistency of the single-cell spatial RNA profiles obtained from images. We argue that it should be possible to match the spatial profiles obtained from images to the scRNA-seq data, and that the percentage of cells that can be matched is thus a quality metric for the segmentation method.

### Reference clusters from scRNA-seq for mouse lung tissue

We re-cluster our single-cell data (32) using only the 6 mapped genes and cells from the same condition (i.e 5 months after irradiation with 17 gy). The final clustering is composed of five different clusters.

### Reference clusters from scRNA-seq for embryonic lung tissue

In the original study (24), the authors applied pciSeq to perform single-cell spatial RNA profiling. Their final expression matrix provided in the study displays 89 genes over the 147 genes map in the HyBISS data. We also keep the same subset of 89 genes for evaluation. We use the same reference scRNA-seq dataset and clustering annotation as provided by the authors.

### Matching in-situ single-cell RNA profile and scRNA-seq

As for simulation, we normalize both scRNA-seq data and the count expression matrix from spatially resolved RNA profiling data with scTransform (37). Many methods exist to match single-cell spatial transcriptomic data and scRNA-seq (38). We chose to leverage the cosine distance as it is robust with respect to different capture rates among the two modalities.

Each cell expression vector was matched to the closest median centroid cluster from scRNA-seq. The cosine distance acts as a proxy for the matching quality.

## Results

### ComSeg overview

Here we present ComSeg, a graph-based method to perform cell segmentation from spatial RNA profiling data. The method operates directly on RNA point clouds and can also leverage cellular landmarks --such as nuclear staining-- when available. For this, we define a KNN graph, where each RNA molecule is a node and where edges are weighted according to the co-expression score of the corresponding genes. Instead of relying on external data to compute these co-expression weights, we leverage the input images by estimating the local gene expression vector in a close environment of each RNA molecule (see Material and Methods). We then detect groups of RNA molecules with similar gene expression in their local environment, by the use of a modified version of the Louvain community detection method (26). Our community detection method can leverage spatial landmarks like DAPI segmentation, as prior knowledge. Lastly, we group the detected RNA communities with similar expression profiles and obtain a transcriptomic domain map at the tissue scale. To obtain single-cell RNA profiles we exploit the cell or nuclei centroid position concurrently to the transcriptomic domain map. An overview of the method is displayed in Figure 1A.

**Figure 1:**
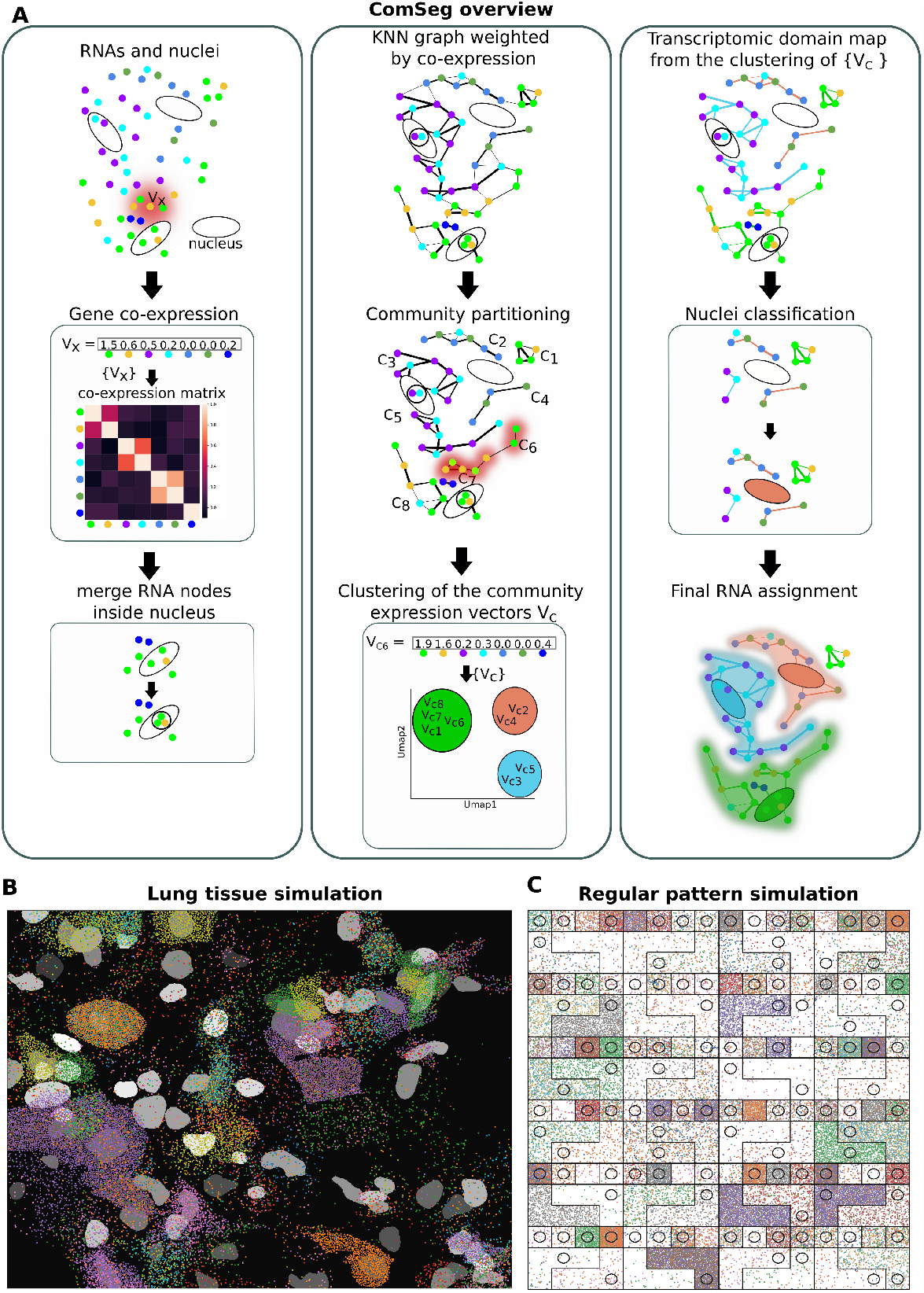
Overview of the method and simulations. **(A)** Overview of the ComSeg method. **(B)** Examples of 3D lung tissue simulation and **(C)** 2D regular pattern simulation with L-shaped cells, both with 34 marker genes represented by different colors.

### State-of-the-art methods for cell segmentation

We benchmarked ComSeg against methods that can be used in an equivalent setting i.e single-cell spatial RNA profiling approaches requiring no external dataset. As a first baseline, we calculated the Watershed transformation (39) as this method is often used for RNA-nuclei association in tissue (40, 41). This method effectively calculates a Voronoi tessellation with the nuclear regions as markers, and is thus equivalent to assigning RNA transcripts to their nearest nucleus. The method hence perfectly works on convex cell shape, if all nuclei are detected, but may fail otherwise. Second, we benchmarked Baysor (18), a cell-marker-free segmentation method optimizing the joint likelihood of transcriptional composition and prior cell morphology. It is particularly suited for cases where only nuclei staining or weak cytoplasmic staining are available. Baysor uses an elliptic function as cell shape prior. To complete this benchmark we also apply a method leveraging external scRNA-seq data, pciSeq (16), a Bayesian model leveraging prior scRNA-Seq data to estimate a probability of cell assignment for each read. Given the observed RNA spot configuration, the method performs cell assignment to match known transcriptomic profiles from scRNA-seq. Hence, it also simultaneously assigns each cell to a cell type. Leveraging prior knowledge may help pciSeq to avoid wrong RNA-cell assignment. However pciSeq implicitly uses a spherical cell shape prior that may hinder its application on complex tissue. Baysor and pciSeq both match their cell shape prior to the RNA spatial distribution to maximize a likelihood function. However, this strategy may fail on sparse data. As sparse RNA point clouds are not a reliable proxy for cell shape, these methods may not identify cell matching their cell shape prior. Hence, on sparse data, these methods might underestimate cell sizes, leading to a high percentage of missing RNAs per cell. The hyper-parameter setting of the benchmarked methods can be found in the Supplementary Methods.

### Benchmark on simulation of lung tissue

To benchmark the methods on challenging data, we turned to simulation mimicking lung tissue (Material and Methods). Here, cells have complex shapes in 3D and airways add empty space devoid of any transcripts. Moreover, we also simulated some cells without nuclei, which can occur in tissue sections. Further, we sampled real expression profiles from our recent scRNA-seq data (32). We simulated 34 cell type marker genes selected with the NS-forest algorithm (36). This marker list enabled us to classify 19 different cell types present in lung tissues with an accuracy of 0.88 for cell type calling when having a perfect RNA assignment to cells (Material and Methods).

To quantitatively compare the different methods, we implemented different metrics (See Material and Methods for details). We assessed the quality of the RNA-cell assignment with the mean Jaccard index per cell, which is calculated for the RNA positions.

The ultimate goal of all the methods is to obtain for each cell a gene expression vector. In this context, not all types of RNA-cell association errors are equivalent. If a cell misses some RNAs, it may still be possible to accurately assess its RNA profile, similarly to cell type classification in scRNAseq, where only a fraction of all RNAs are sequenced. However, if some RNAs are wrongly associated with a cell, this can create a mixed expression profile resulting in wrong cell type classification. We thus reported the mean percentage of wrongly associated RNA per cell (*WA*) and the mean percentage of missing RNA per cell (*MS*) separately. Lastly, we assessed cell type calling accuracy by comparison to the ground truth cell type defined by scRNA-seq data (Material and Methods).

Examples of RNA-cell assignment from the different approaches on lung tissue simulation are shown in Figure 2A. ComSeg outperforms Baysor, pciSeq and the Watershed algorithm with a Jaccard Index of 0.57 against 0.30, 0.33 and 0.50 for Baysor, pciSeq and Watershed, respectively (Figure 2B, left panel). A closer look at the results, reveals that the type of error is not the same for the four models. Baysor and pciSeq have a high percentage (more than 50%) of missing RNA per cell (Figure 2C, right panel) while Watershed has a very low (15%) mean percentage of missing RNA per cell. This is not surprising, as the Watershed computes a full Voronoi Tessellation and hence assigns all RNAs. Watershed and pciSeq have a higher mean percentage of wrongly associated RNA per cell (about 40%) while it is low for Baysor and Comseg (roughly 20% on average) (Figure 2C, left panel).

**Figure 2:**
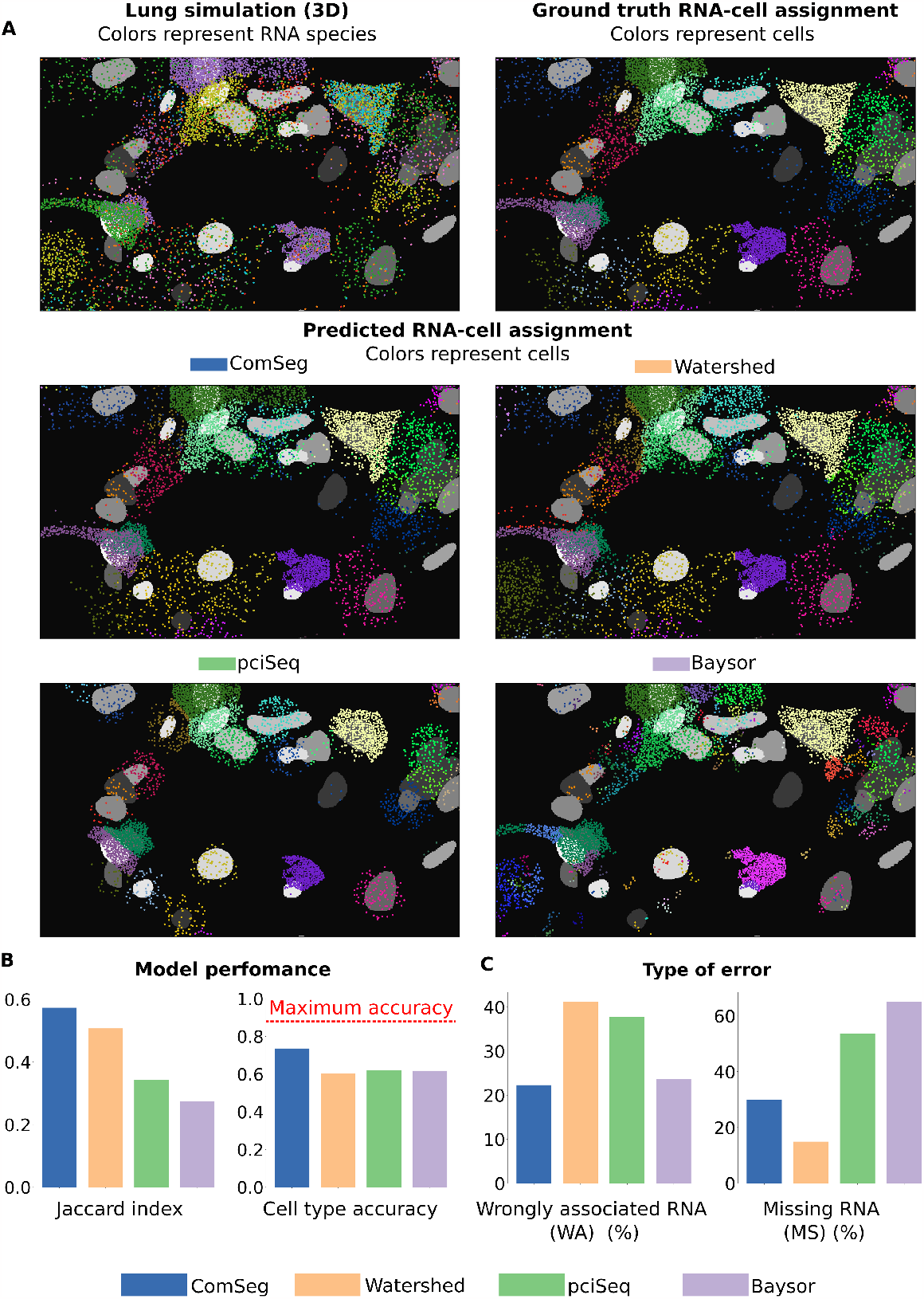
Benchmarking on simulated lung tissue.(**A**)3D lung simulation and RNA assignment of the different models. The ground truth takes into account only cells with a nucleus.(**B**)Mean Jaccard in per cell (left panel) and cell type calling accuracy (right panel) for the benchmarked models, the red line is the maximum accuracy when having a perfect RNA assignment to cells.(**C**)Mean percentage of wrongly associated RNA per cell (*WA*) (left panel) and mean percentage of missing RNA per cell (*MS*) (right panel).

Next, we compared the estimated expression profiles from the simulated images to the known, underlying scRNA-seq ground-truth, by performing cell type calling. Here, ComSeg reaches 74%, compared to 60%, 62% and 62% for Watershed, pciSeq and Baysor respectively (Figure 2B, right panel). The higher accuracy of ComSeg for cell type calling can be explained by the lower misassociation error rate (Figure 2C, left panel). Of note, the best cell type calling accuracy that could be achieved is 88%, which is thus the value that would be reached if all RNAs were correctly assigned to the cell they belong to. This value is not 100%, because the selected marker genes do not perfectly recapitulate the full transcriptomic profile. On the other hand, we observe that cell type calling is heavily impacted by wrong RNA assignments.

In conclusion, ComSeg performs better RNA-cell association and significantly improves downstream tasks like cell type calling on lung tissue simulation.

In view of these results, we next turned to simulations with a simpler tissue geometry, in order to better understand the limitations of each of the methods.

### Validation on simple simulations of regular patterns

In this section we aim to better understand the strengths and limitations of each benchmarked method. To this end, for validation and benchmarking, we generated five types of simulated datasets of gradually increasing complexity and specific cases.

To study the effect of the different possible expression profiles without any cell shape complexity, we simulate a checkerboard (see Material and Methods). This is a similar cell size scale to what we can observe in mouse tissue (32, 42). To test the effect of non-convex shapes, we also simulate L-shaped cells (see Material and Methods). Each simulated set contains 10 images of 110 cells except the last one containing 144 cells per image.

#### 1) Simulation 1 (Sim 1). Variation of expression level

The objective of this simulation is to benchmark the methods when markers have the same expression level versus the case where one marker is sparsely expressed. We started by simulating only two cell types expressing one marker each, A and B. We fixed the number of transcripts for the first cell type to A=100, while we set the number of transcripts for the second to B=100 (Sim1a) or B=10 (Sim1b) in two runs (Figure 3A, left panels). For this geometry, the Watershed performs a perfect assignment, as the nuclei are centered and the cell shapes are convex. Besides, in the simple case where the two markers are equally expressed, Baysor and ComSeg have also a Jaccard index close to 1 whereas pciSeq has a Jaccard index below 0.8. In fact, pciSeq only assigns RNAs in the close neighborhood of the nucleus and with a spherical shape prior. These missed RNAs are penalized by the Jaccard index.

**Figure 3:**
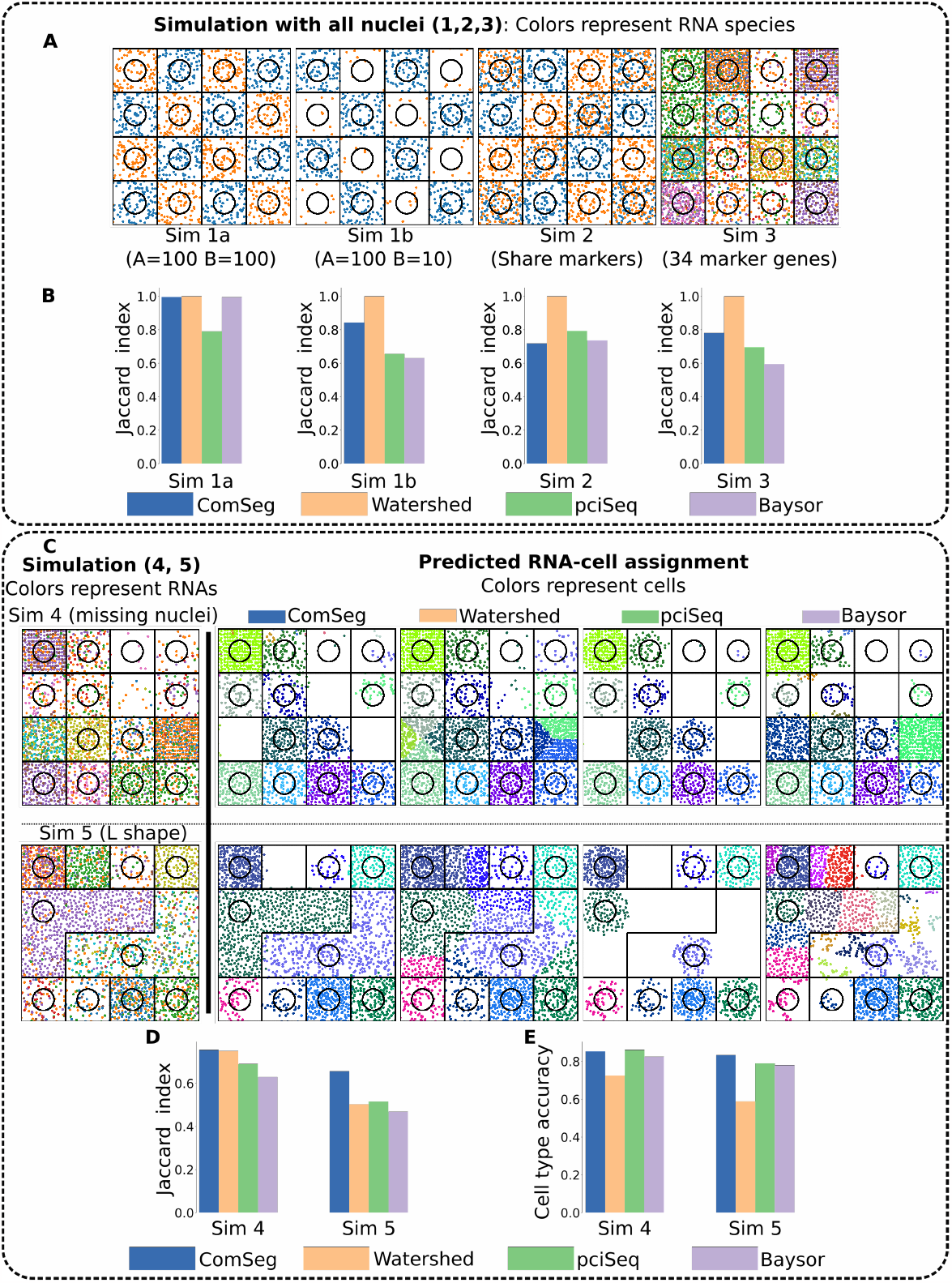
Simulations and results on regular patterns. **(A)** Examples of simulation where all cells have a nucleus: Sim 1 with variation of expression level, Sim 2 with share marker genes, Sim 3 with experimental RNA profiles with 34 markers. **(B)**

When the expression of the second cell type becomes sparse (B=10), the performance of all methods leveraging RNA spatial distribution, Baysor, pciSeq and ComSeg drops. Still, ComSeg performs better in terms of Jaccard index (over 0.8,Figure 3B, center left panel) and cell type accuracy (of 0.99, Supplementary Figure 1) than Baysor and pciSeq. This result can be attributed to the utilization of a cell shape prior by Baysor and pciSeq in RNA assignment. This approach proves less effective when expression becomes sparse, as Baysor and pciSeq may not find RNA point cloud matching their shape prior. Finally, the shape agnostic strategy of ComSeg appears to be more adapted for sparse input.

#### 2) Simulation 2 (Sim 2). Shared marker genes

In the subsequent simulation, we aimed to investigate the impact of shared marker genes across diverse cell types. To achieve this, we categorized three distinct cell types: cell type A, with 100 RNAs from gene A; cell type B, with 100 RNAs from gene B; and cell type C with 100 RNAs from gene A and gene B (Figure 3A, center right panel). In this case pciSeq and Baysor have slightly better performance in terms of Jaccard index than ComSeg (Figure 3B, center right panel). Still, all models get an almost perfect cell type calling (Supplementary Figure 1). Without surprise, Watershed obtains perfect RNA-nuclei assignment due to the simplicity of the tissue geometry.

#### 3) Simulation 3 (Sim 3). Experimental expression profile

In this simulation, we sample RNA profiles from experimental data. We simulate 34 marker genes as described above for lung tissue simulation (Figure 3A, right panel). On real RNA profiles, ComSeg has a better Jaccard index than Baysor and pciSeq. As the cell shapes are still convex, Watershed gets a Jaccard index of 1 (Figure 3B, right panel).

#### 4) Simulation 4 (Sim 4). Experimental expression profile with missing nuclei

In this simulation, a scenario akin to the previous one was replicated, where some nuclei were intentionally omitted to simulate conditions akin to experimental data (Figure 3C, left panel, first row). Specifically, 20% of the cells were simulated without nuclei, mirroring situations encountered in tissue experiments. Remarkably, under these conditions, ComSeg continued to exhibit a superior Jaccard index compared to both Baysor and pciSeq (Figure 3D, left). As expected, the Watershed method cannot cope with missing nuclei, as the method uses nuclei as seeds. As a consequence, accuracy in cell type identification drops as compared to other models. (Figure 3E, left panel). A visual representation of the benchmarked methods RNA assignments can be found in Figure 3C, first row.

3)Simulation 5 (Sim 5). Experimental expression profile with missing nuclei and L shape:

Lastly we also add cells with an L-shape to test non-convex examples (Figure 3C, left panel, second row). In this case, ComSeg outperforms other models for the Jaccard index confirming that it is designed to deal with irregular non-convex cell shape. Conversely, Watershed has a very low Jaccard index because it cannot cope with non-convex shapes by construction (Figure 3D, right panel). Also, Baysor and pciSeq are underperforming in accurately identifying the L-shaped cells owing to their inherently convex cell shape assumptions. As a consequence, ComSeg also exhibits superior cell type calling performances in this more complex case as compared to all other methods (Figure 3E, right panel). A visual representation of the RNA assignments generated by all models can be found in Figure 3C, second row.

In summary, all methodologies encounter difficulties as cell shapes and RNA profiles become increasingly complex and as marker expression becomes sparse. Notably, Watershed proves to be an optimal choice for cells with convex shapes. PciSeq and Baysor, while capable of estimating valid single-cell spatial RNA profiles in terms of cell type calling, exhibit limitations in capturing a substantial portion of transcripts. Moreover, the disparity in RNA-cell assignment performance between ComSeg and the other benchmarked methods widens notably as cell shapes and expression patterns grow in complexity.

Mean Jaccard index per cell for Sim 1-2-3.(C, left)Example of Sim 4 with experimental RNA profiles with 34 markers and missing nucleus and Sim 5 akin to Sim 4 but with L-shaped cells and(C, right)the corresponding RNA-cell assignment for the benchmarked methods. (D)Mean Jaccard index per cell and(E)cell type calling accuracy for Sim 4 and Sim 5.

### Application to experimental data

We applied ComSeg to two different experimental lung datasets, each with a specific challenge for the analysis. The first dataset was created in-house. In this experiment, we visualized 6 different marker genes in 3D mouse lung tissue. Our approach has a very high RNA detection efficiency, but many cells display no RNA transcripts as only a subset of cell types is targeted. The second dataset is from a recent study of human embryonic lung mapping 147 genes in 2D (24). The HybISS approach used for this dataset has a lower capture rate (15) but enables the visualization of many more genes.

For such experimental data, no ground truth is available. In tissue with complex morphology such has lung, having no ground truth makes it particularly challenging to assess the method’s quality. An existing validation strategy is based on the correlation of segmented regions from different methods (15, 43). However, this validation method compares methods between themselves. Errors in existing methods could thus be propagated to the next generation of methods. In the absence of direct access to ground truth for imaging datasets, we chose to leverage available scRNA-seq datasets obtained from identical organ samples. These scRNA-seq datasets serve as a means to assess the consistency of single-cell spatial RNA profiling. As for simulations, we calculate the cosine distance between cell expression vectors derived from images and the nearest scRNA-seq cluster centroid. Consequently, the cosine distance between single-cell RNA profiles from the image dataset and scRNA-seq clusters serves as a surrogate measure for the quality of single-cell spatial RNA profiling.

Similarly to what we did on simulations, we applied Baysor, pciSeq, Watershed and ComSeg on the 3D mouse lung tissue dataset (Figure 4A) and on the 2D embryonic lung tissue dataset.

**Figure 4:**
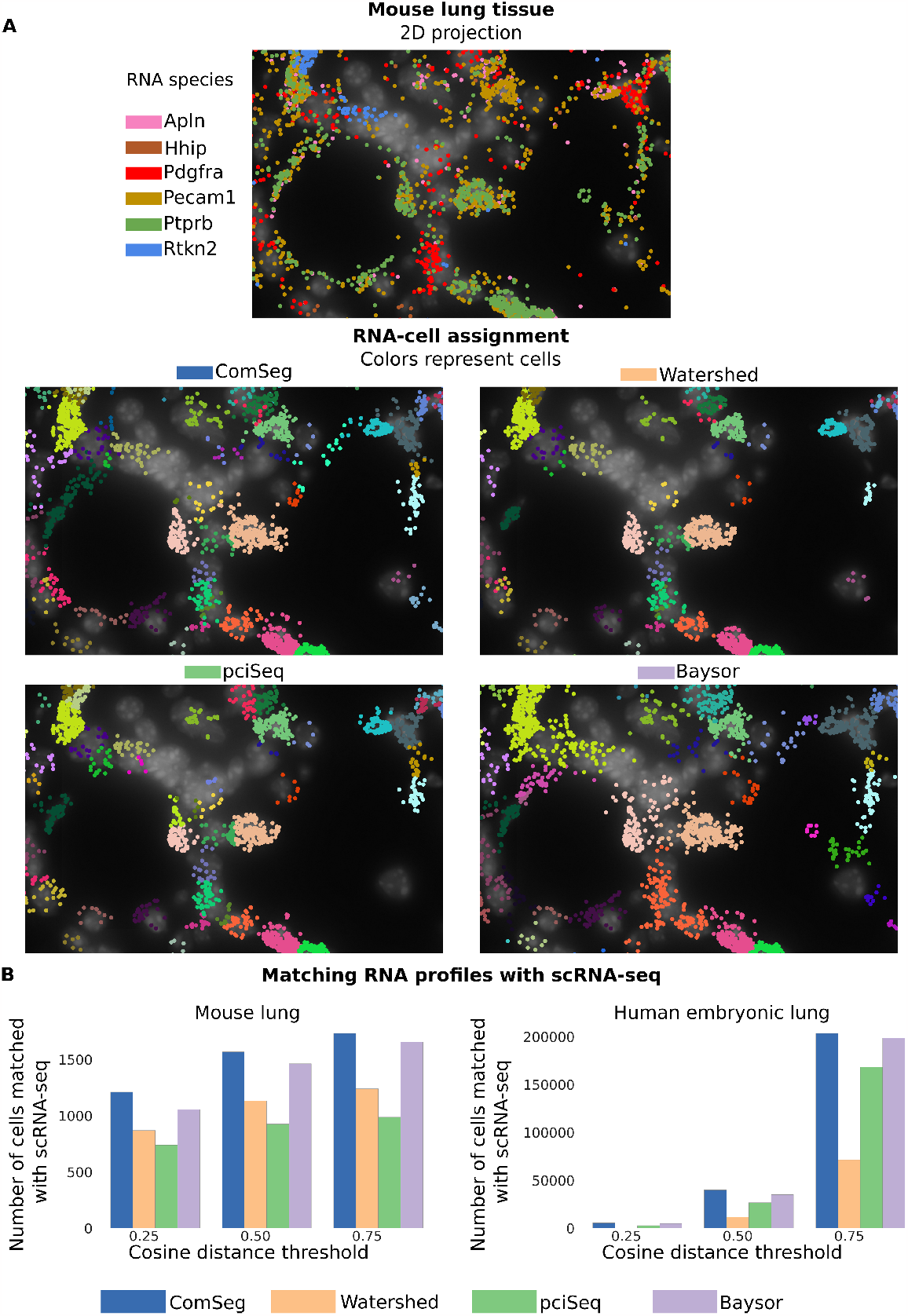
Benchmark on experimental data.(A)Mouse lung tissue and RNA assignment of the benchmarked models.(B)Number of RNA profiles identified in images matched with a scRNA-seq reference dataset at different cosine distance thresholds for mouse lung tissue and human embryonic lung tissue.

On mouse lung tissue, ComSeg identifies more cells than the other tested methods, for which an RNA profile can be assigned. (Supplementary Figure 2, left). We obtain similar results for human embryonic lung tissue (Supplementary Figure 2, right) where ComSeg also detects more cells than the three other methods we tested. We took into account only cells with more than 5 RNAs. Importantly, the number of cells with matching expression profiles in the scRNA-seq data is higher for ComSeg for both mouse (Figure 4B, right panel) and human lung tissue (Figure 4B, left panel) at the different cosine distance threshold.

In summary, ComSeg detects more cells than other methods, and our analysis revealed that the RNA profiles measured in the segmented cells also fit better external datasets, thus suggesting better segmentation quality. These experimental results confirm the results of the simulation: ComSeg is suitable for complex cell shapes, as observed in lung tissue.

## Discussion

Imaging-based spatial RNA profiling methods provide RNA point cloud coordinates without information about the cell origin of each molecule. Still, it is essential to correctly assign RNAs to cells to perform downstream tasks at the single-cell level such as cell type calling or inference of cell interaction. In this study, we present a new method called ComSeg, a graph-based method operating directly on the RNA coordinates. ComSeg can leverage landmark images such as nuclear staining. In addition, ComSeg is cell shape agnostic which makes it particularly useful for complex tissues, where cell shape convexity cannot be assumed. As such, ComSeg is a flexible method able to handle various situations, demonstrating good performances even in challenging experimental settings.

In order to compare ComSeg to other methods, we have developed a simulation framework (*SimTissue*) that provides tissue architectures which are generated from real images and are therefore reasonably realistic. Our simulation environment also encompasses simple geometric shapes which are ideally suited to study failure modes and limitations of algorithms thanks to the simplified geometry. Indeed, quality assessment is a critical aspect for segmentation, but actually difficult to perform. So far, there are no manually annotated ground truth for imaging-based spatial RNA profiling. Comparison to a consensus is an option that has been adopted by several authors (15, 43). We argue that this can lead to propagation of systematic errors. For instance, all competing methods rely on shape priors that are not always met in complex tissues. A comparison to the consensus would not be able to reveal this.

We found that most methods have difficulties with non-convex shapes as well as large differences in gene expression density. Our simulations suggest that error rates in cell type calling are non-negligible: as much as 26% of cell type assignments are erroneous because of segmentation errors when using previously published methods. ComSeg outperforms competing methods by a large margin on complex tissues. However, even with ComSeg, cell classification errors due to wrong segmentations amount to 14%, thus suggesting that development of novel segmentation methods remains an important topic for imaging-based spatial RNA profiling data.

Beyond these benchmark results, ComSeg has several other compelling aspects that makes it interesting for the scientific community. First, it does not require cellular landmarks, such as membrane stainings. Indeed, existing membrane stainings are highly variable and not very robust. Moreover, they are inhomogeneous and can therefore lead to spatial biases in cell type calling accuracies. Second, ComSeg does not require external datasets, such as scRNAseq, which makes it also applicable in small-scale studies, where such data is not available. Lastly, ComSeg does not rely on annotated data, which is very tedious and sometimes impossible to provide. Its modular structure makes it easy to tailor ComSeg to particularities in the datasets.

One of the limitations of ComSeg is that it is dependent on the choice of cell type marker genes. The model may fail if the spatially resolved RNA species are not discriminative of cell type or cell state. A potential improvement of ComSeg would be to incorporate in the model several landmark stainings.

In all, we believe that this model will be of great interest to the community and has the potential to overcome current shortcomings in cell segmentation and cell type calling from spatial RNA profiling data. To facilitate the use of ComSeg, we have made it available as an open-source and documented Python package: https://github.com/tdefa/ComSeg. We also make the simulation framework S*imTissue*(https://github.com/tdefa/SimTissue) available to the community, which might help researchers in the future to benchmark their methods.

## Data availibity

The implementation code of ComSeg is available at https://github.com/tdefa/ComSeg_pkg. The implementation code of SimTissue is available at https://github.com/tdefa/SimTissue. The simulated dataset of lung tissue can be downloaded from Zenodo (https://zenodo.org/records/10172316). Scripts and datasets to reproduce the benchmark are available also on Zenodo at https://zenodo.org/records/10222671.

## Acknowledgements

This work has received financial support through the Agence Nationale de la Recherche (ANR) grant (LUSTRA, reference ANR-19-CE14-0015-04 to A.L.-V., C.F., F.Mu. and T.W.). F.Mu. and C.W. acknowledge funding by Institut Pasteur. Furthermore, this work was supported by the French government under management of Agence Nationale de la Recherche as part of the ‘‘Investissements d’avenir’’ program, reference ANR-19-P3IA-0001 (PRAIRIE 3IA Institute).

S.C.A. and A.L.-V. also received support from La Ligue Contre Le cancer. S.C.A. and A.M. were recipients of PhD fellowships from the European Union’s Horizon 2020 research and innovation program under the Marie Skłodowska-Curie grant agreement No 666003 and No 847718 respectively. H.L. is the recipient of a PhD fellowship from the International Student program from Paris-Saclay University. J.S. is recipient of a PhD fellowship from the French Ministry of Education, research and Industry.

## Supplementary Method

### Hyper-parameter setting of benchmarked methods

#### ComSeg

We apply ComSeg with mean cell diameter *D*= 10μ*m* and *R*_*max*_= 15μ*m* in checkerboard simulation (Sim 1-2-3-4) and increase to *R*_*max*_ = 40μ *m* in Sim 5 while keeping the same value for *D*.

We set *D*= 15μ *m* and *R*_*max*_ = 30μ *m* in lung tissue simulation. We use *D*= 20μ *m* and *R*_*max*_ = 40μ *m* on mouse lung tissue and *D*= 40*pixels* and *D*= 50 *pixels* on embryonic lung tissue.

#### Baysor

We apply Baysor by incorporating nucleus segmentation mask priors and let Baysor estimate the scale parameter automatically. All other parameters were left as default.

Baysor does not perform RNA-nucleus assignment but groups RNAs into cells without referring to the given nucleus segmentation index of the prior segmentation mask. On simulated datasets, in order to compare Baysor to other methods for RNA-nucleus assignment, we associated each predicted cell index by Baysor with a nucleus segmentation index from the provided segmentation mask ground truth. To that end, each predicted cell index by Baysor was associated with the nucleus index from the ground truth with the most molecules in common.

#### pciSeq

We apply pciSeq on geometric simulation using simulated scRNA-seq data containing the simulated RNA profiles. In lung tissue simulations, we use the same scRNA-seq dataset as the one sampled to simulate RNA profiles. All other parameters were left as default. We use the version 0.0.46 of the pciSeq Python PyPI package. We use maximum projection to apply pciSeq to 3D data as pciSeq is only designed for 2D.

#### Watershed

We apply Watershed by taking as input the inverse distance map from segmented nuclei. In lung simulation, as a mask, we use the inferred cytoplasm from the Cy3 signal. Otherwise, we use the Watershed on the inverse distance map with a maximum distance from the nucleus of 2 μm on mouse lung tissue and of 16 pixels for embryonic lung tissue. The mask helps to prevent the misassignment of RNA to nuclei too far appart.

### Influence of initialization seed on ComSeg

ComSeg employs random initialization within the Louvain method when computing the communities and the Leiden method when clustering the {*V*_*C*_ }. To evaluate the impact of the random initialization, we executed the algorithm 50 times on the lung simulation and compared the mean Jaccard index per cell for each run with a different random initialization (Supplementary Figure 3). The standard deviation of the mean Jaccard index per cell is less than 0.002, demonstrating the negligible influence of the initialization seed.

## Supplementary figures

**Supplementary Figure 1:**
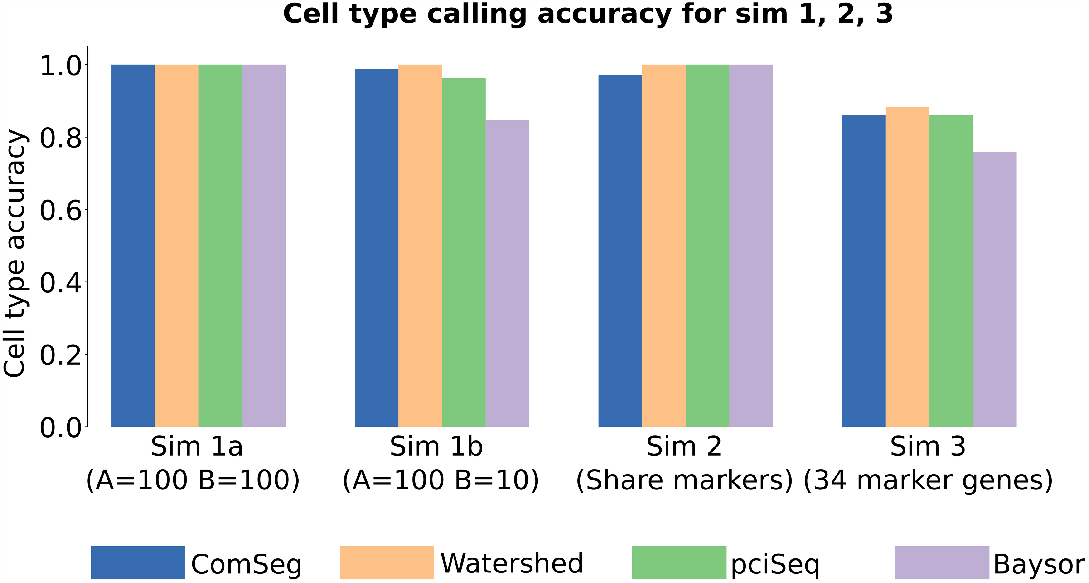
Cell type calling accuracy on checkerboard cell for the Sim 1 with different levels of expressions, Sim 2 with share markers and sim 3 with 34 marker genes with RNA profiles sample from scRNA-seq.

**Supplementary Figure 2:**
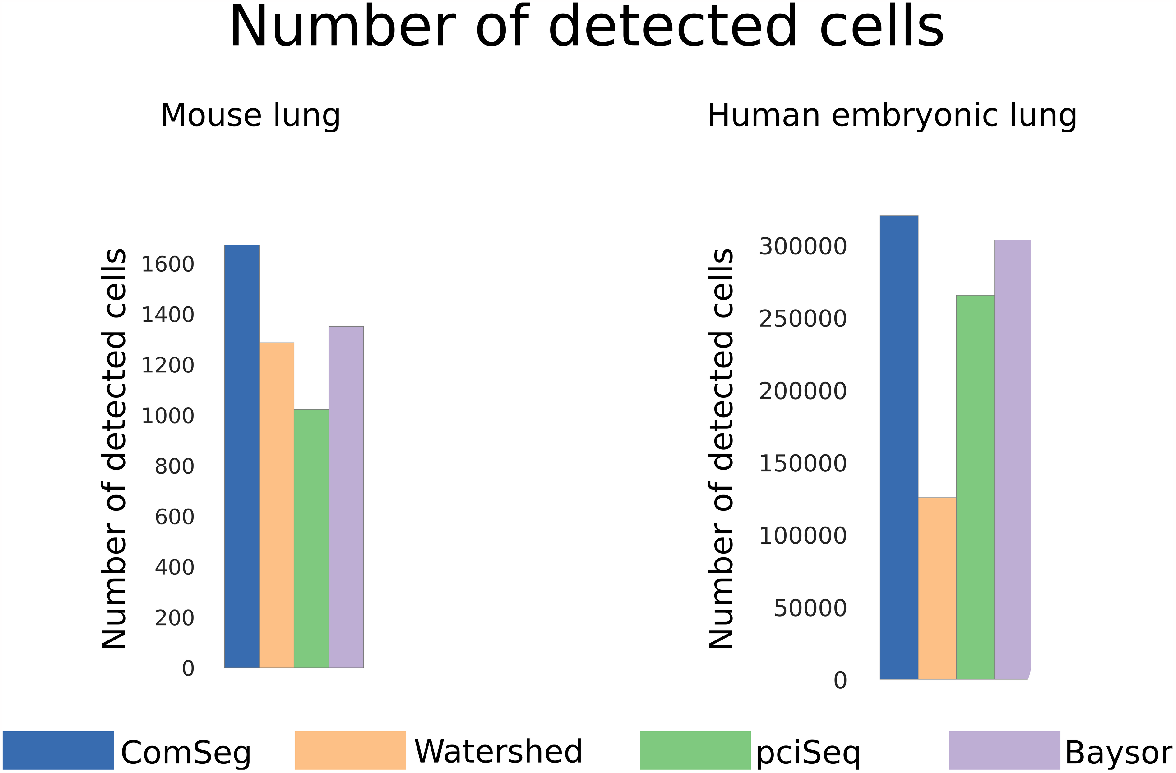
Number of cells associated with an RNA profile (more than 5 RNAs) in the mouse lung tissue dataset (right) and in the Human embryonic lung tissue dataset (Left) for the tested method.

**Supplementary Figure 3:**
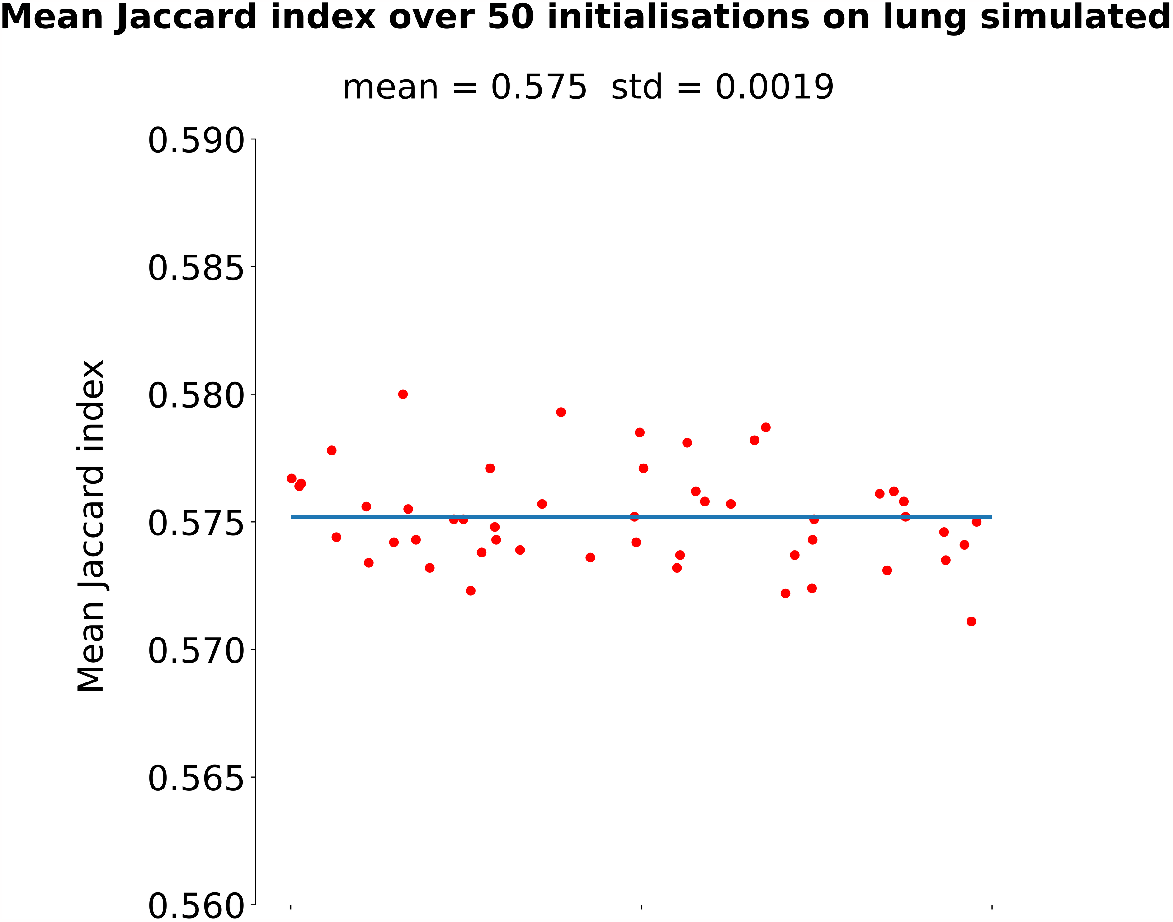
The mean Jaccard index per cell obtained for 50 different random initializations of ComSeg on our simulated dataset of lung tissue. The blue line is the mean of all the Jaccard index obtained for 50 different random initializations.

## Notes

### Competing Interest Statement

The authors have declared no competing interest.

